# ProtSpace: a tool for visualizing protein space

**DOI:** 10.1101/2024.11.30.626168

**Authors:** Tobias Senoner, Tobias Olenyi, Michael Heinzinger, Anton Spannagl, George Bouras, Burkhard Rost, Ivan Koludarov

**Author notes:** Corresponding author: Tobias Senoner. Senior authors contributed equally to this work□ http://www.rostlab.org, (email rost) (Burkhard Rost, Ivan Koludarov).

## Abstract

Protein language models (pLMs) generate high-dimensional representations of proteins, so called embeddings, that capture complex information stored in the set of evolved sequences. Interpreting these embeddings remains an important challenge. *ProtSpace* provides one solution through an open-source Python package that visualizes protein embeddings interactively in 2D and 3D. The combination of embedding space with protein 3D structure view aids in discovering functional patterns readily missed by traditional sequence analysis.

We present two examples to showcase *ProtSpace*. First, investigations of phage data sets showed distinct clusters of major functional groups and a mixed region, possibly suggesting bias in today’s protein sequences used to train pLMs. Second, the analysis of venom proteins revealed unexpected convergent evolution between scorpion and snake toxins; this challenges existing toxin family classifications and added evidence refuting the *aculeatoxin family hypothesis*.

*ProtSpace* is freely available as a pip-installable Python package (source code & documentation) with examples on GitHub (https://github.com/tsenoner/protspace) and as a web interface (https://protspace.rostlab.org). The platform enables seamless collaboration through portable JSON session files.

## Introduction

The concept of “protein space”, introduced by Maynard Smith in 1970 [1], provides a powerful framework for understanding protein evolution: a vast multidimensional landscape in which each point represents a unique sequence surrounded by direct neighbors differing by single mutations. This abstract space, combined with Wright’s concept of fitness landscapes [2] applied to proteins [3], helps explain how proteins navigate evolutionary trajectories through sequence changes that maintain or alter biological function. While traditionally explored through sequence similarity metrics, protein Language Models (pLMs) such as ProtTrans [4], ESM3 [5], and ProtGPT2 [6] have revolutionized our ability to represent this space through high-dimensional protein embeddings: these learned representations encode amino acid sequences into numerical vectors that capture non-local structural and functional relations between proteins [7]. Embeddings from pLMs increasingly replacing traditional multiple sequence alignments (MSAs) as input features for many residue- and protein-level prediction tasks, from secondary structure prediction to protein function classification [8–13].

Although pLMs are transforming computational biology, their high-dimensional embeddings present a significant challenge: unlike traditional MSAs, these complex representations are not directly interpretable. This creates a barrier, particularly for experts who want to leverage their domain knowledge in conjunction with the representation when generating hypotheses. *ProtSpace* addresses this challenge; an open-source Python package makes protein embeddings accessible through interactive 2D and 3D visualizations where researchers can explore the relationship between embeddings and their expert-curated annotations. Our tool uniquely integrates protein-specific features, including 3D structure visualization, while enabling the creation and sharing of analysis sessions.

Earlier visualization approaches for protein relationships include CLANS [14], which enables exploration of all-against-all BLAST sequence similarities through force-directed graph layout. The EFI-EST web tool [15] further expanded this concept by streamlining the generation of sequence similarity networks with automated processing of protein families and enhanced visualization capabilities. More general embedding visualization tools, while not specifically designed for proteins, include Google’s Embedding Projector [16], Parallax [17], and Whatlies [18], which offer interactive exploration of high-dimensional embeddings but lack protein-specific features. The ESM Metagenomic Atlas [19] provides a specialized platform for exploring pre-computed ESM2 embeddings and their structures predicted with ESMFold for metagenomic sequences, though it is limited to preloaded datasets and does not allow users to explore their own.

Recent evolutionary studies demonstrated the potential of *ProtSpace* by revealing functional clustering patterns in snake three-finger toxins by visualizing embedding space [20] and by helping to disprove the existence of a hypothetical aculeatoxin protein family [21] in *Hymenoptera* venoms [22]. Here, we present an extended, detailed analysis of *ProtSpace* visualizations and demonstrate how researchers can leverage this tool for their own datasets through our freely available web service (https://protspace.rostlab.org) and Python package (https://github.com/tsenoner/protspace).

## Materials and methods

### Data collection

We curated two distinct protein datasets to demonstrate different applications of protein embedding visualization. Each dataset was processed through our standardized pipeline (described below) while addressing domain-specific requirements. The datasets and relevant splits are available on GitHub (https://github.com/tsenoner/protspace/tree/main/data).

#### Toxin Dataset

We collected venom protein sequences from SwissProt [23] (release 2024_02) using the keyword KW-0800 (“toxin”) and filtered for venom tissue specificity in Metazoa. Signal peptides were predicted and removed using SignalP6.0 [24] to focus the analysis on mature peptide sequences. Each sequence was annotated with taxonomic classification and functional information from UniProt, including protein family membership. The final dataset comprised 5,181 sequences spanning 18 taxonomic orders and 28 protein families.

#### Phages Dataset

The PHROG dataset is a library of viral protein families constructed through remote homology detection via HMM profile-profile comparisons, providing deeper clustering than BLAST-like similarity searches while maintaining functional homogeneity [25]. Each protein family contains a higher-level annotation from 9 categories describing phage biology (e.g., tail, head and packaging, or lysis) and a specific annotation (e.g., tail fiber protein, major capsid protein, or endolysin). From 938,864 proteins in the complete database, we analyzed 7,500 proteins. We first identified 370,000 Foldseek [26] cluster representatives (at 30% sequence identity threshold) and generated protein structures using Colabfold v1.5.5 [27]. The final dataset comprised 7,000 functionally annotated proteins spanning all 9 categories and 500 hypothetical proteins, ensuring both structural and sequence diversity in the selected subset.

### Data Preparation

*ProtSpace* transforms protein sequence data into interactive visualizations through a structured preprocessing workflow. The complete pipeline is illustrated in **Fig. S1** and detailed in our Jupyter notebooks (available in our GitHub repository). It consists of three main steps: embedding generation, dimensionality reduction, and annotation integration.

The workflow supports two input types: pLM embeddings or similarity matrices. For embedding generation, we employed ProtT5-XL-U50 (hereafter ProtT5 [4]) accessed via the HuggingFace model repository [28]. For a protein of length L, the model outputs a 1024-dimensional vector for each residue, resulting in an L x 1024 matrix. We mean-pool these vectors across residues to obtain a fixed-length per-protein embedding. These high-dimensional embeddings are then projected into visualizable 2D or 3D coordinates using established dimensionality reduction techniques: Principal component analysis (PCA) [29], Multidimensional scaling (MDS) [30], or Uniform Manifold Approximation and Projection (UMAP) [31]. Alternatively, users can provide similarity matrices (e.g. sequence or structure similarity) instead of pLM-derived embeddings; they will be processed and displayed accordingly.

The final step combines the reduced embeddings with user-provided annotations of interest, such as functional classifications, experimental measurements, or taxonomic information. All information is consolidated into a standardized JSON format that serves as input for *ProtSpace* visualization and analysis.

### ProtSpace implementation

The *ProtSpace* dashboard was implemented using Dash and Plotly [32], enabling responsive data exploration. 3D protein structure visualization was integrated using NGL Viewer [33]. *ProtSpace* is accessible as a web service (https://protspace.rostlab.org/) and can be installed locally as a Python package. The tool supports exporting 2D plots as SVG files, 3D plots as interactive HTML files, and complete workspaces as JSON files for reproducibility and collaboration.

## Results and discussion

### ProtSpace revealed functional organization and pLM limitations in phage proteomes

*ProtSpace* visualization of phage protein embeddings revealed clear functional organization with several unexpected patterns (**Figure 1**). Major functional components clustered distinctly in embedding space: structural elements formed two separate clusters at the right and left of the space, while proteins involved in nucleotide metabolism occupied the central region. Notably, lysis-related proteins formed three distinct clusters, suggesting potential functional sub-specialization in cell lysis mechanisms.

**Figure 1:**
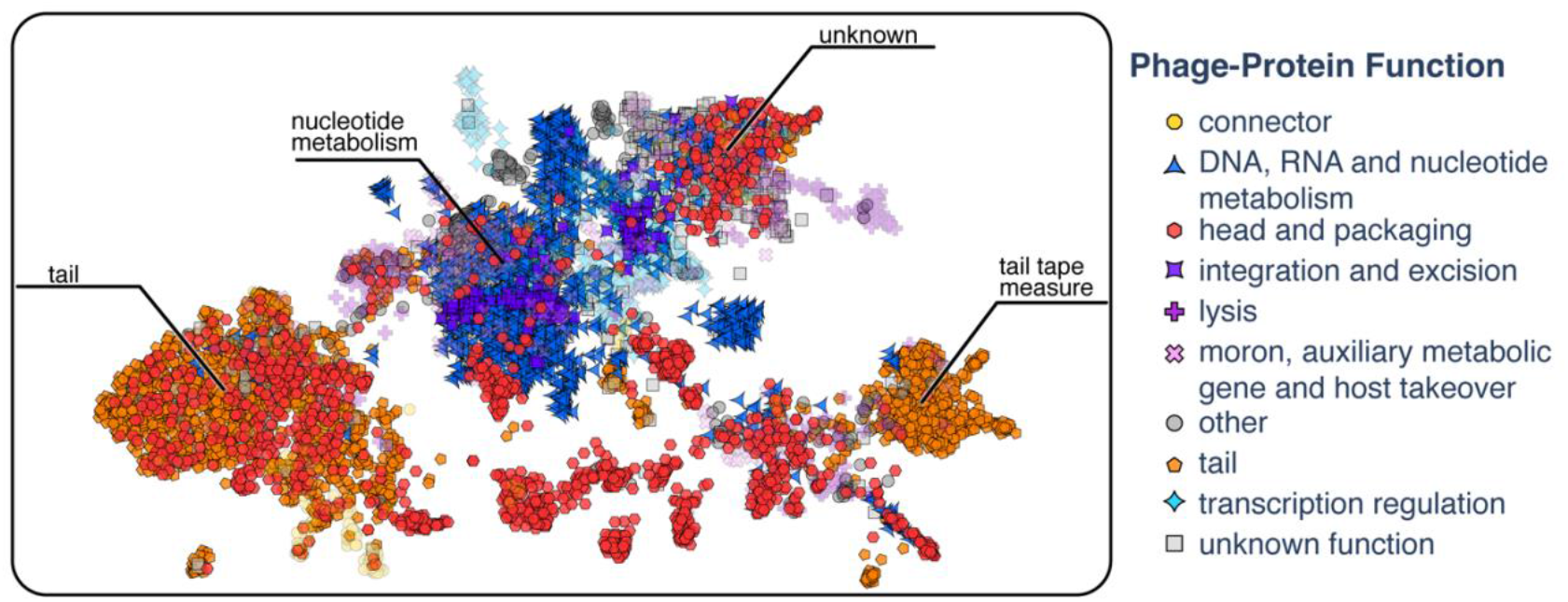
Protein embeddings distinguish major structural protein elements in Phages. Two-dimensional UMAP projection of ProtT5 [4] embeddings, where each point represents a phage protein. Major structural components analyzed in the manuscript are shown in vibrant colors, while other protein groups appear desaturated for easier visualization. The complete visualization with more detailed protein categories can be found in **Fig. S2**.

Finer-grained analysis of structural proteins (**Fig. S2**) revealed specific organization within these clusters. The right structural cluster contained primarily tail tape measure proteins, while the left cluster combined both major and minor tail proteins along with other structural elements. This clear separation of tail tape measure proteins suggests distinct sequence patterns for this specialized structural component.

Most intriguingly, *ProtSpace* also pointed to limitations in the pLM training. A region in the top contained proteins with known, diverse functions that nevertheless clustered together. Moreover, this region showed a mixed distribution of various functional categories, suggesting these sequences might be outside the model’s training distribution. This observation generates a testable hypothesis about the relationship between model training data and phage protein diversity.

### Protein Language Models reveal evolutionary convergence and divergence in animal toxins

The venom dataset represents a highly curated subset of UniProt [23] with protein family nomenclature based on previous independent sequence and structure similarity studies. Analysis of this dataset in *ProtSpace* revealed distinct taxonomic and functional clusters (**Figure 2** and **Fig. S3**). Toxins of marine snails (Neogastropoda, mostly *Conus*) and arachnids (Araneae, mostly spiders) occupy the central space, while aculeatan (Hymenoptera, including ants, wasps, and bees) and caterpillar (Lepidoptera) proteins cluster in the bottom-right center region. Toxins from scorpions and centipedes blend from the right and left toward the middle, while several distinct reptilian (Squamata, primarily snakes) and arachnid clusters appear separate to the periphery, suggesting independent evolution of unique toxin families. These clusters all correspond to major venom protein families, including three-finger toxins, phospholipases, metalloproteinases, and snake c-type lectins.

**Figure 2:**
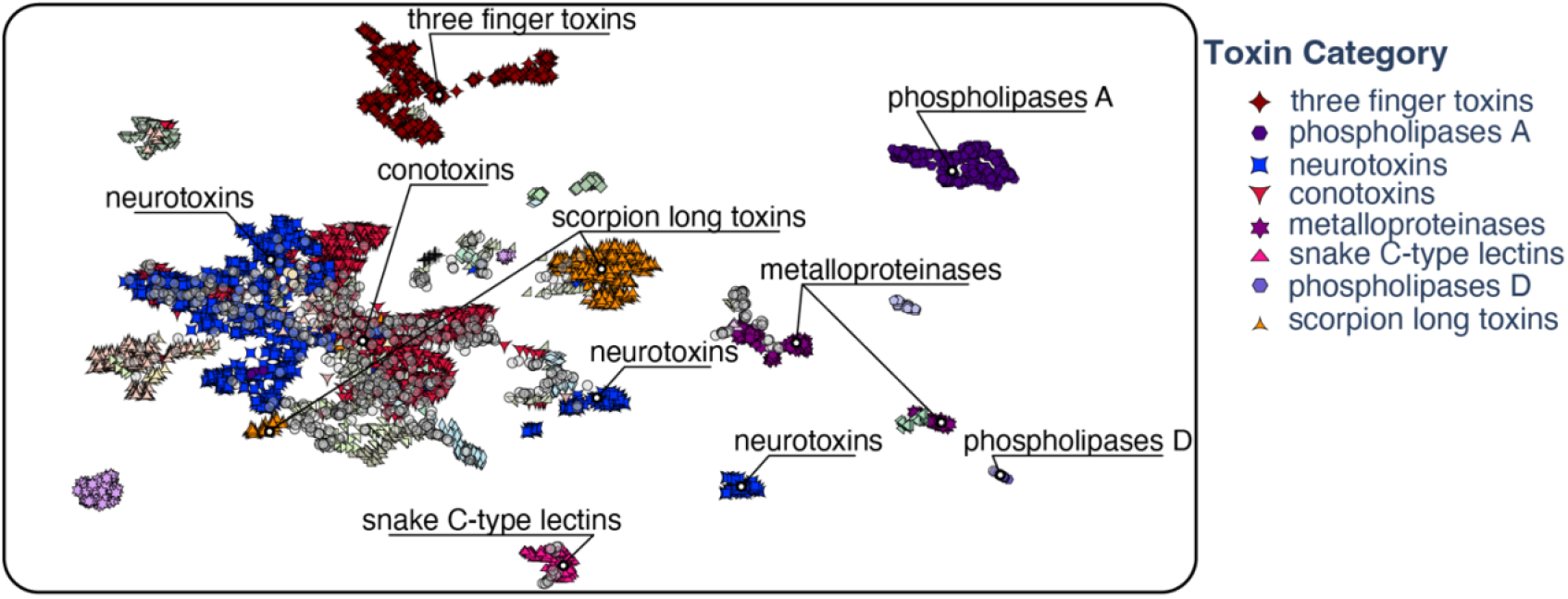
pLM embeddings visualized in 2D reveal functional clustering of animal toxins. Two-dimensional UMAP projection of ProtT5 [4] embedding vectors, where each point represents a venom protein. Points are colored and shaped according to different protein families. Protein families discussed in the manuscript are shown in vibrant colors, while other families appear desaturated. The original image can be found in **Fig. S3**.

The *ProtSpace* analysis reveals inconsistencies in current toxin nomenclature. Specifically, proteins labeled as “neurotoxin” split into three distinct clusters suggesting potential functional subdivisions within these families of predominantly spider toxins. Similarly, the “scorpion long toxin” family forms two separate clusters, likely reflecting a propensity to bind different ion channels. Notably, seven centipede toxins (UniProt IDs: I6R1R5, P0DPX5, P0DPX9, P0DPY0, P0DPY1, P0DPX7, P0DPX8) cluster with snake three-finger toxins despite distinct taxonomic, and likely evolutionary origins. Structural investigation of these unique centipede toxins revealed remarkable similarity to the three-finger toxin fold, suggesting either convergent evolution between scorpion and snake proteins or descent from a common distant ancestor – both representing remarkable possibilities warranting a more thorough study.

Another interesting find is that mast-cell degranulating family of Hymenopteran toxins separates into two distinct clusters that align with the split between bee apamin-related and wasp mastoparan-related peptides, supporting previous genomics-based results [22] indicating these represent independent protein lineages achieving similar functions through convergent evolution, thus invalidating their classification as a single protein family.

Further functional divergence appears in metalloproteinases, where disintegrin and non-disintegrin proteins form separate clusters, further supporting their classification as distinct protein lineages. Phospholipases A and D occupy distinct regions of the embedding space. The Phospholipase A cluster comprises reptilian, scorpion, starfish, and fang blenny fish representatives, while Phospholipase D contains arachnid, one scorpion, and one cone snail protein. The latter cluster warrants deeper investigation, potentially representing an independently evolved lineage unrelated to canonical phospholipases.

These patterns revealed through embedding space visualization, represent only the most prominent protein relationships in the dataset yet generate testable hypotheses about toxin evolution and challenge existing nomenclature and classification systems derived solely from sequence similarity or taxonomic relationships.

## Conclusions

*ProtSpace* reveals the power and limitations of embeddings from protein Language Model (pLMs) embeddings through interactive visualization. The tool identified how phages harbor out-of-distribution protein sequences for ProtT5 and exposed previously unknown evolutionary convergence in toxin families. A key innovation of *ProtSpace* is its integration of structural and embedding space representations. The tool efficiently handles datasets of up to 20,000 proteins while maintaining responsive interactivity, features one-click generation of quality figures, and allows complete analysis sessions to be shared through single JSON files. These capabilities have already enabled new insights into protein evolution, as demonstrated by our mapping of snake three-finger toxins protein space to show how secreted forms evolved from membrane-anchored ones, and our in-depth analysis of Hymenopteran (wasps, bees, ants) venom proteins that helped to refute a hypothesis about a hypothetical aculeatoxin protein family.

## Abbreviations

2D: two-dimensional
3D: three-dimensional
BLAST: Basic Local Alignment Search Tool
CLANS: CLuster ANalysis of Sequences
EFI-EST: Enzyme Function Initiative-Enzyme Similarity Tool
HTML: Hypertext Markup Language (file format)
HMM: Hidden Markov Model
JSON: JavaScript Object Notation (file format)
MDS: Multidimensional scaling
MSA: multiple sequence alignments
PCA: Principal component analysis
PHROG: protein orthologous groups
pLM: protein Language Model
SVG: Scalable Vector Graphics (file format)
UMAP: Uniform Manifold Approximation and Projection.

## Declarations

## Data availability

The complete source code, documentation, and example datasets are available through GitHub (https://github.com/tsenoner/protspace) under the AGPL-3.0 license. All datasets used in the analyses are stored in the repository’s data directory (https://github.com/tsenoner/protspace/tree/main/data). The interactive visualization platform is freely accessible at https://protspace.rostlab.org/.

## Competing interests

The authors declare no competing interests.

## Authors’ contributions

Tobias Senoner: Conceptualization, Data curation, Methodology, Software, Visualization, Writing – original draft. Tobias Olenyi: Resources, Software, Writing – review & editing. Michael Heinzinger: Conceptualization, Software, Writing – review & editing. Anton Spannagle: Software. George Bouras: Resources, Writing – review & editing. Burkhard Rost: Supervision, Scientific oversight, Writing – review & editing. Ivan Koludarov: Conceptualization, Data curation, Visualization, Supervision, Writing – original draft.

## Funding

The Bavarian Ministry of Education supported the work through funding to the TUM. The work was also supported by the German Ministry for Research and Education (BMBF: Bundesministerium für Bildung und Forschung); BMBF Projekt 16DKWN13, SSTDBB.

## Acknowledgments

We would like to thank Nikita Kugut for his support with many aspects of this work. We extend our gratitude to the Technical University of Munich (TUM) for providing the necessary facilities and resources for this research. Additionally, we are grateful to all contributors to open-source programming libraries. Finally, we thank those who deposit experimental data in public databases, maintain these databases, and develop methods to enrich experimental data.

## Notes

### Competing Interest Statement

The authors have declared no competing interest.

## References

[1] J. Maynard Smith, Natural Selection and the Concept of a Protein Space, Nature 225 (1970) 563–564. 10.1038/225563a0.

[2] S. Wright, The Roles of Mutation, Inbreeding, crossbreeding and Selection in Evolution, Proceedings of the XI International Congress of Genetics 8 (1932) 209–222.

[3] J.A.G.M. De Visser, J. Krug, Empirical fitness landscapes and the predictability of evolution, Nat Rev Genet 15 (2014) 480–490. 10.1038/nrg3744.

[4] A. Elnaggar, M. Heinzinger, C. Dallago, G. Rehawi, Y. Wang, L. Jones, T. Gibbs, T. Feher, C. Angerer, M. Steinegger, D. Bhowmik, B. Rost, ProtTrans: Toward Understanding the Language of Life Through Self-Supervised Learning, IEEE Trans. Pattern Anal. Mach. Intell. 44 (2022) 7112–7127. 10.1109/TPAMI.2021.3095381.

[5] T. Hayes, R. Rao, H. Akin, N.J. Sofroniew, D. Oktay, Z. Lin, R. Verkuil, V.Q. Tran, J. Deaton, M. Wiggert, R. Badkundri, I. Shafkat, J. Gong, A. Derry, R.S. Molina, N. Thomas, Y. Khan, C. Mishra, C. Kim, L.J. Bartie, M. Nemeth, P.D. Hsu, T. Sercu, S. Candido, A. Rives, Simulating 500 million years of evolution with a language model, (2024). 10.1101/2024.07.01.600583.

[6] N. Ferruz, S. Schmidt, B. Höcker, ProtGPT2 is a deep unsupervised language model for protein design, Nat Commun 13 (2022) 4348. 10.1038/s41467-022-32007-7.

[7] M. Littmann, M. Heinzinger, C. Dallago, T. Olenyi, B. Rost, Embeddings from deep learning transfer GO annotations beyond homology, Sci Rep 11 (2021) 1160. 10.1038/s41598-020-80786-0.

[8] M. Bernhofer, B. Rost, TMbed: transmembrane proteins predicted through language model embeddings, BMC Bioinformatics 23 (2022) 326. 10.1186/s12859-022-04873-x.

[9] M.L. Bileschi, D. Belanger, D.H. Bryant, T. Sanderson, B. Carter, D. Sculley, A. Bateman, M.A. DePristo, L.J. Colwell, Using deep learning to annotate the protein universe, Nat Biotechnol 40 (2022) 932–937. 10.1038/s41587-021-01179-w.

[10] D. Ilzhöfer, M. Heinzinger, B. Rost, SETH predicts nuances of residue disorder from protein embeddings, Front. Bioinform. 2 (2022) 1019597. 10.3389/fbinf.2022.1019597.

[11] J. Meier, R. Rao, R. Verkuil, J. Liu, T. Sercu, A. Rives, Language models enable zero-shot prediction of the effects of mutations on protein function, (2021). 10.1101/2021.07.09.450648.

[12] H. Stärk, C. Dallago, M. Heinzinger, B. Rost, Light attention predicts protein location from the language of life, Bioinformatics Advances 1 (2021) vbab035. 10.1093/bioadv/vbab035.

[13] M. Manfredi, C. Savojardo, P.L. Martelli, R. Casadio, ISPRED-SEQ: Deep Neural Networks and Embeddings for Predicting Interaction Sites in Protein Sequences, Journal of Molecular Biology 435 (2023) 167963. 10.1016/j.jmb.2023.167963.

[14] T. Frickey, A. Lupas, CLANS: a Java application for visualizing protein families based on pairwise similarity, Bioinformatics 20 (2004) 3702–3704. 10.1093/bioinformatics/bth444.

[15] J.A. Gerlt, J.T. Bouvier, D.B. Davidson, H.J. Imker, B. Sadkhin, D.R. Slater, K.L. Whalen, Enzyme Function Initiative-Enzyme Similarity Tool (EFI-EST): A web tool for generating protein sequence similarity networks, Biochimica et Biophysica Acta (BBA) - Proteins and Proteomics 1854 (2015) 1019–1037. 10.1016/j.bbapap.2015.04.015.

[16] D. Smilkov, N. Thorat, C. Nicholson, E. Reif, F.B. Viégas, M. Wattenberg, Embedding Projector: Interactive Visualization and Interpretation of Embeddings, (2016). http://arxiv.org/abs/1611.05469 (accessed February 28, 2023).

[17] P. Molino, Y. Wang, J. Zhang, Parallax: Visualizing and Understanding the Semantics of Embedding Spaces via Algebraic Formulae, in: Proceedings of the 57th Annual Meeting of the Association for Computational Linguistics: System Demonstrations, Association for Computational Linguistics, Florence, Italy, 2019: pp. 165–180. 10.18653/v1/P19-3028.

[18] V. Warmerdam, T. Kober, R. Tatman, Going Beyond T-SNE: Exposing whatlies in Text Embeddings, in: Proceedings of Second Workshop for NLP Open Source Software (NLP-OSS), Association for Computational Linguistics, Online, 2020: pp. 52–60. 10.18653/v1/2020.nlposs-1.8.

[19] Z. Lin, H. Akin, R. Rao, B. Hie, Z. Zhu, W. Lu, N. Smetanin, R. Verkuil, O. Kabeli, Y. Shmueli, M. Fazel-Zarandi, T. Sercu, S. Candido, A. Rives, Evolutionary-scale prediction of atomic-level protein structure with a language model, (2023).

[20] I. Koludarov, M. Heinzinger, T.N.W. Jackson, D. Dashevsky, S.D. Aird, B. Rost, Domain loss enables evolution of novel functions in a gene superfamily, including snake 3-finger toxins, Molecular Biology, 2022. 10.1101/2022.12.15.520616.

[21] S.D. Robinson, A. Mueller, D. Clayton, H. Starobova, B.R. Hamilton, R.J. Payne, I. Vetter, G.F. King, E.A.B. Undheim, A comprehensive portrait of the venom of the giant red bull ant, Myrmecia gulosa, reveals a hyperdiverse hymenopteran toxin gene family, SCIENCE ADVANCES (2018).

[22] I. Koludarov, M. Velasque, T. Senoner, T. Timm, C. Greve, A.B. Hamadou, D.K. Gupta, G. Lochnit, M. Heinzinger, A. Vilcinskas, R. Gloag, B.A. Harpur, L. Podsiadlowski, B. Rost, T.N.W. Jackson, S. Dutertre, E. Stolle, B.M. Von Reumont, Prevalent bee venom genes evolved before the aculeate stinger and eusociality, BMC Biol 21 (2023) 229. 10.1186/s12915-023-01656-5.

[23] The UniProt Consortium, A. Bateman, M.-J. Martin, S. Orchard, M. Magrane, S. Ahmad, E. Alpi, E.H. Bowler-Barnett, R. Britto, H. Bye-A-Jee, A. Cukura, P. Denny, T. Dogan, T. Ebenezer, J. Fan, P. Garmiri, L.J. Da Costa Gonzales, E. Hatton-Ellis, A. Hussein, A. Ignatchenko, G. Insana, R. Ishtiaq, V. Joshi, D. Jyothi, S. Kandasaamy, A. Lock, A. Luciani, M. Lugaric, J. Luo, Y. Lussi, A. MacDougall, F. Madeira, M. Mahmoudy, A. Mishra, K. Moulang, A. Nightingale, S. Pundir, G. Qi, S. Raj, P. Raposo, D.L. Rice, R. Saidi, R. Santos, E. Speretta, J. Stephenson, P. Totoo, E. Turner, N. Tyagi, P. Vasudev, K. Warner, X. Watkins, R. Zaru, H. Zellner, A.J. Bridge, L. Aimo, G. Argoud-Puy, A.H. Auchincloss, K.B. Axelsen, P. Bansal, D. Baratin, T.M. Batista Neto, M.-C. Blatter, J.T. Bolleman, E. Boutet, L. Breuza, B.C. Gil, C. Casals-Casas, K.C. Echioukh, E. Coudert, B. Cuche, E. De Castro, A. Estreicher, M.L. Famiglietti, M. Feuermann, E. Gasteiger, P. Gaudet, S. Gehant, V. Gerritsen, A. Gos, N. Gruaz, C. Hulo, N. Hyka-Nouspikel, F. Jungo, A. Kerhornou, P. Le Mercier, D. Lieberherr, P. Masson, A. Morgat, V. Muthukrishnan, S. Paesano, I. Pedruzzi, S. Pilbout, L. Pourcel, S. Poux, M. Pozzato, M. Pruess, N. Redaschi, C. Rivoire, C.J.A. Sigrist, K. Sonesson, S. Sundaram, C.H. Wu, C.N. Arighi, L. Arminski, C. Chen, Y. Chen, H. Huang, K. Laiho, P. McGarvey, D.A. Natale, K. Ross, C.R. Vinayaka, Q. Wang, Y. Wang, J. Zhang, UniProt: the Universal Protein Knowledgebase in 2023, Nucleic Acids Research 51 (2023) D523–D531. 10.1093/nar/gkac1052.

[24] F. Teufel, J.J. Almagro Armenteros, A.R. Johansen, M.H. Gíslason, S.I. Pihl, K.D. Tsirigos, O. Winther, S. Brunak, G. Von Heijne, H. Nielsen, SignalP 6.0 predicts all five types of signal peptides using protein language models, Nat Biotechnol 40 (2022) 1023–1025. 10.1038/s41587-021-01156-3.

[25] P. Terzian, E. Olo Ndela, C. Galiez, J. Lossouarn, R.E. Pérez Bucio, R. Mom, A. Toussaint, M.-A. Petit, F. Enault, PHROG: families of prokaryotic virus proteins clustered using remote homology, NAR Genomics and Bioinformatics 3 (2021) qab067. 10.1093/nargab/lqab067.

[26] M. Van Kempen, S.S. Kim, C. Tumescheit, M. Mirdita, J. Lee, C.L.M. Gilchrist, J. Söding, M. Steinegger, Fast and accurate protein structure search with Foldseek, Nat Biotechnol 42 (2024) 243–246. 10.1038/s41587-023-01773-0.

[27] M. Mirdita, K. Schütze, Y. Moriwaki, L. Heo, S. Ovchinnikov, M. Steinegger, ColabFold: making protein folding accessible to all, Nat Methods 19 (2022) 679–682. 10.1038/s41592-022-01488-1.

[28] T. Wolf, L. Debut, V. Sanh, J. Chaumond, C. Delangue, A. Moi, P. Cistac, T. Rault, R. Louf, M. Funtowicz, J. Davison, S. Shleifer, P. Von Platen, C. Ma, Y. Jernite, J. Plu, C. Xu, T. Le Scao, S. Gugger, M. Drame, Q. Lhoest, A. Rush, Transformers: State-of-the-Art Natural Language Processing, in: Proceedings of the 2020 Conference on Empirical Methods in Natural Language Processing: System Demonstrations, Association for Computational Linguistics, Online, 2020: pp. 38–45. 10.18653/v1/2020.emnlp-demos.6.

[29] K. Pearson, LIII. On lines and planes of closest fit to systems of points in space, The London, Edinburgh, and Dublin Philosophical Magazine and Journal of Science 2 (1901) 559–572. 10.1080/14786440109462720.

[30] W.S. Torgerson, Multidimensional scaling: I. Theory and method, Psychometrika 17 (1952) 401–419. 10.1007/BF02288916.

[31] L. McInnes, J. Healy, J. Melville, UMAP: Uniform Manifold Approximation and Projection for Dimension Reduction, (2020). http://arxiv.org/abs/1802.03426 (accessed October 3, 2024).

[32] S. Hossain, Visualization of Bioinformatics Data with Dash Bio, in: C. Calloway, D. Lippa, D. Niederhut, D. Shupe (Eds.), Proceedings of the 18th Python in Science Conference, 2019: pp. 126–133. 10.25080/Majora-7ddc1dd1-012.

[33] A.S. Rose, P.W. Hildebrand, NGL Viewer: a web application for molecular visualization, Nucleic Acids Res 43 (2015) W576–W579. 10.1093/nar/gkv402.

