## supplementary-material for "ProtSpace: a tool for visualizing protein space"

### ProtSpace: a tool for visualizing protein space - Supporting Material

[1]Technical University of Munich (TUM)), School of Computation, Information and Technology (CIT), Faculty of Informatics, Chair of Bioinformatics & Computational Biology, [2]Adelaide Medical School, Faculty of Health and Medical Sciences, The University of Adelaide, [3]The Department of Surgery—Otolaryngology Head and Neck Surgery, University of Adelaide and the Basil Hetzel Institute for Translational Health Research, Central Adelaide Local Health Network, [4]Institute for Advanced Study (TUM-IAS), [5]TUM School of Life Sciences Weihenstephan

#### Short Description

This supplementary document provides additional visualizations and analyses to complement the main manuscript. We present a workflow overview of ProtSpace (**Fig. S1**). Additionally, we provide detailed extended figures from the:

- Phages dataset (**Fig. S2**)
- Toxin dataset (**Fig. S3**)

#### Table of Contents for SOM

**Submitted** Nov 30, 2024

#### Figure S1: ProtSpace Workflow overview

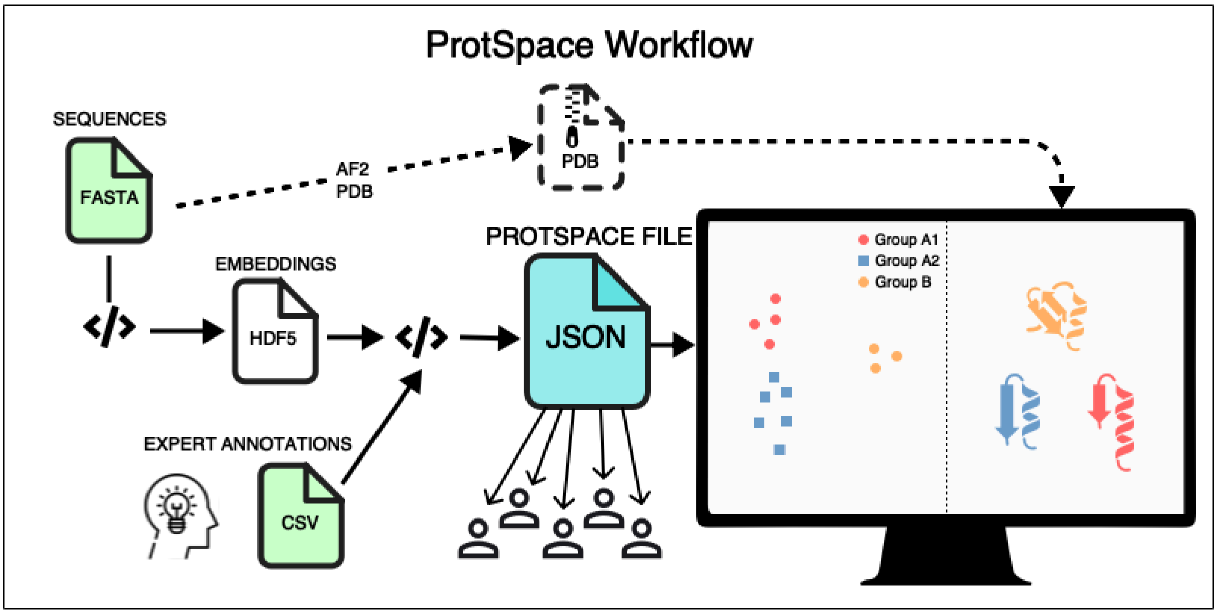

**Fig. S1**: **ProtSpace integrates embeddings, expert knowledge, and structure for protein space exploration.** Overview of ProtSpace which transforms protein sequences and expert knowledge (input files with green background) into an interactive visualization. Protein sequences from a FASTA file are converted into per-protein embeddings and stored in an HDF5 file. These embeddings, combined with user-defined labels of interest, are processed by a preparation script to generate a JSON file (output file with turquoise background). This JSON file serves as the input for the ProtSpace visualization tool and can be shared with colleagues, facilitating the collaborative exploration of the protein space. For structural analysis, users can optionally provide a ZIP file containing PDB files of protein structures.

#### Figure S2: 2D visualization of phage protein classifications

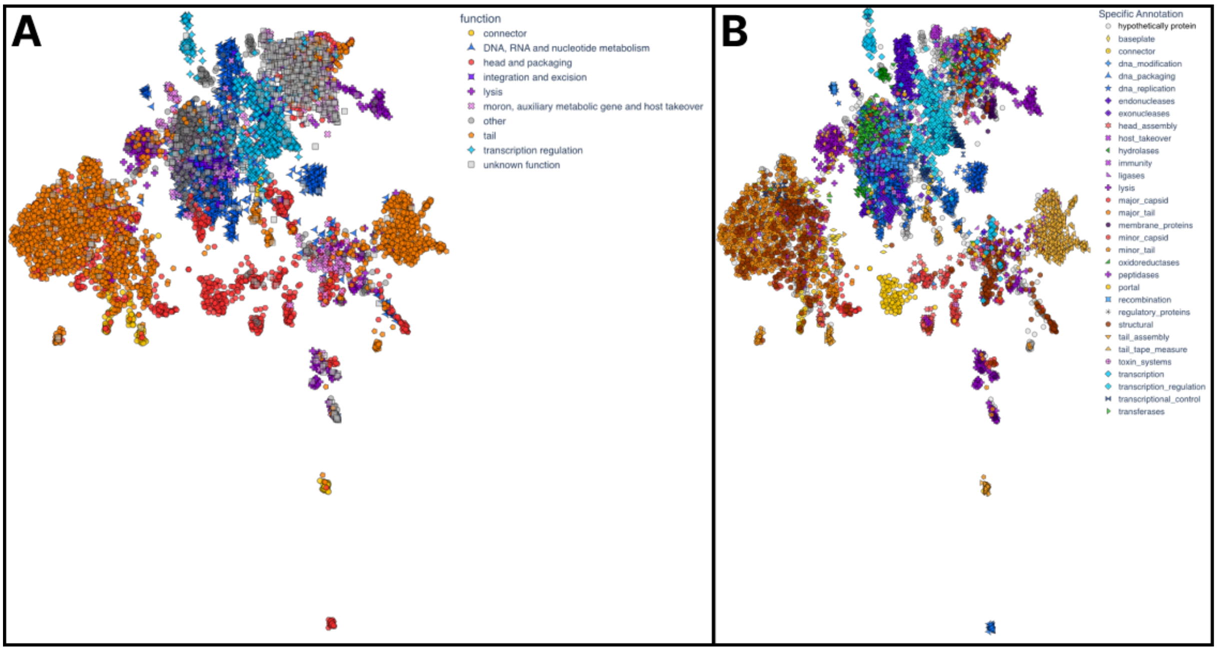

**Fig. S2**: **Comparative visualization of coarse and fine-grained phage protein classifications** Two-dimensional UMAP projections of ProtT5 embeddings for phage proteins. Panel A) shows the grouping of 9 major functional categories as presented in Figure 1 without color editing. Panel B) displays the same spatial arrangement with detailed product subcategories, revealing finer granularity in structural and functional protein clustering.

#### Figure S3: 2D visualization of venom protein families

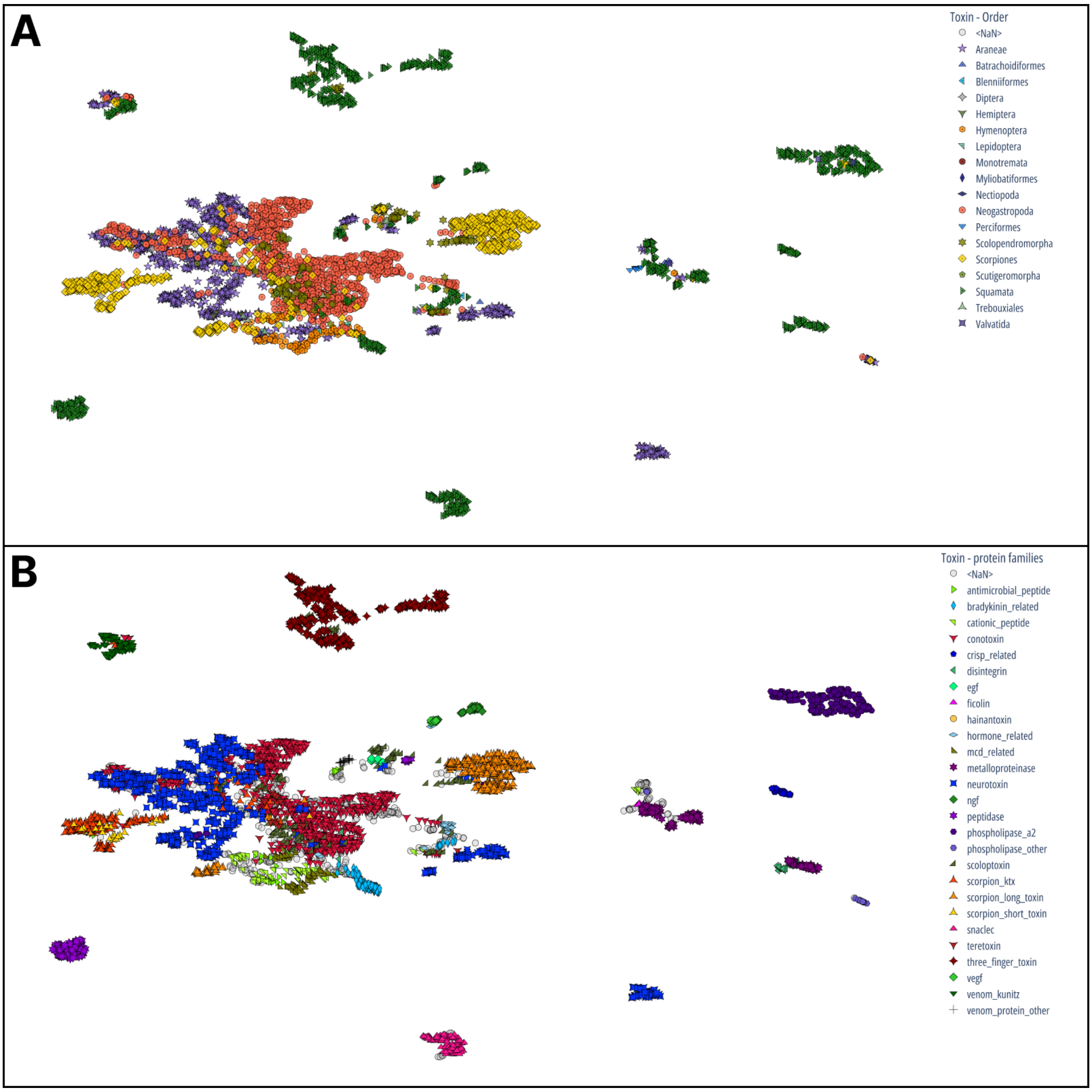

**Fig. S3**: **Comparative UMAP visualization of protein categories and taxonomic orders in venom protein embeddings** Side-by-side UMAP projections of the same protein language model embedding vectors. Panel A) shows proteins colored by taxonomic order. Panel B) displays the same spatial arrangement colored and shaped by protein category as in Figure 1, but with all major families' labels shown. This comparison allows direct visualization of how protein functions are distributed across taxonomic groups in the embedding space.
